# Transient and DNA-free *in vivo* CRISPR/Cas9 genome edition for flexible modelling of endometrial carcinogenesis

**DOI:** 10.1101/2022.06.17.496593

**Authors:** Raúl Navaridas, Maria Vidal-Sabanés, Anna Ruiz-Mitjana, Aida Perramon-Güell, Cristina Megino-Luque, David Llobet-Navas, Xavier Matias-Guiu, Joaquim Egea, Mario Encinas, Lídia Bardia, Julien Colombelli, Xavier Dolcet

## Abstract

The CRISPR/Cas9 technology has emerged as a powerful tool to generate mouse models of disease. Endometrial cancer is the most common malignancy of the female genital tract. In the present study, we have developed a pipeline for the generation of somatically engineered mouse models of endometrial cancer by *in vivo* electroporation-mediated delivery of Cas9 ribonucleoprotein into the uterine cavity. By using mT/mG dual-fluorescent reporter mice, we show that this system allows an efficient genomic edition specifically in epithelial endometrial cells. As a proof of its applicability for endometrial cancer modeling, we designed Cas9 ribonucleoprotein targeting Pten, the most frequently tumor suppressor gene mutated in this type of cancer. Pten-targeting ribonucleoprotein delivery into the uterine cavity caused loss of expression of PTEN protein in epithelial endometrial cells that resulted in the development of endometrial lesions. We also validated this technique for gene edition in other important endometrial driver genes such as *p53* or *Fbxw7*. By co-targeting *LoxP* sites of mT/mG reporter mice and Pten, we demonstrate the generation of differentially edited cell populations that may be a useful tool to model tumoral heterogeneity. Moreover, the combination of CRISPR/Cas9 technology in mT/mG dual-reporter mice and light-sheet microscopy represents an interesting approach for *in vivo* cancer cell tracing. This methodology opens a new door for future rapid, flexible, customizable and multiplexable *in vivo* modeling of endometrial cancer.

## INTRODUCTION

The clustered regularly interspaced short palindromic repeats (CRISPR) and CRISPR-associated (Cas) proteins were identified as part of prokaryotic adaptive immune system (1). Among all CRISPR/Cas9 existing systems across the prokaryotes (2), the *Streptococcus pyogenes* (SpCas9) endonuclease was the first and more widely used system for biological applications of genome edition. SpCas9 is an RNA-guided endonuclease that generates double strand-breaks (DSBs) on target DNA sequences with the only requirement of an ‘*NGG*’ protospacer adjacent motif (PAM) on 3’ end of target sequence. Cas9-associated guide RNA (gRNA) directs Cas9 nuclease to complementary target sequences. Since the implementation of CRISPR/Cas9 as a tool for mammalian genome edition (3–6), an overwhelming number of CRISPR/Cas9 systems have been applied to virtually all fields of biomedical sciences (2, 7). In cancer research, the CRISPR/Cas9-mediated genome edition has been a powerful tool to advance in the knowledge of many aspects of cancer such as genetics and epigenetics, cancer cell heterogeneity, or therapeutics (8, 9).

*In vivo* modelling of cancer using genetically engineered mouse models (GEMM) provided unfading information about the role of genes alterations in the development and progression of cancer. Traditionally, generation of either constitutive or conditional germline GEMMs has been the main approach to mimic human cancer-associated molecular alterations in mice (10). In the last decade, CRISPR/Cas9-mediated edition of embryonic stem cells or oocytes has emerged as an efficient and technically improved method to generate germline GEMM of cancer (8). Besides germline generation of knock-out or knock-in mice, CRISPR/Cas9-mediated generation of somatically GEMM have emerged as a new approach for *in vivo* modelling of cancer for basic and translational research (11). Somatic GEMM carrying single or combinations of mutations have been developed to model different neoplasms such as acute myeloid leukemia (12), liver and pancreatic cancer (13–15) or lung adenocarcinomas (16).

For delivery of CRISPR/Cas9 components (Cas9 and gRNA) into embryonic or adult mammalian cells two main choices need to be considered: the type of molecule in which CRISPR/Cas9 components are delivered and the method used to introduce these molecules into the living cells. Cas9 and gRNA can be delivered as plasmid DNA encoding for both Cas9 and gRNA, as a combination of RNAs (mRNA for Cas9+gRNA), or as a ribonucleoprotein (RNP) formed by the recombinant Cas9 and gRNA (17). Introduction of these molecules into the cell can be performed by classical transfection techniques including microinjection, lipofection, or electroporation (18, 19), by viral transduction or by an increasing number of novel materials and strategies (18).

Endometrial cancer (EC) is the most frequent type of cancer in the female reproductive tract (20). Germline GEMMs that target the major drivers of EC have been used to model the pathogenesis of this type of neoplasm (21, 22). In the late 1990s *Pten* was identified as a frequently mutated gene in EC (23–25) and, in the last decade, genomic landscape of EC confirmed *Pten* as the most frequently mutated tumor suppressor in EC (26, 27). The role of *Pten* mutations in the EC development has been confirmed by the generation of genetically engineered mouse models carrying heterozygous deletion of *Pten* (28–30) or, more specifically, by conditional deletion of *Pten* in the uterus or endometrial epithelium (21).

In the present study, we have developed a novel protocol for the generation of somatically GEMM of EC *in vivo*, by intra-uterine electroporation of Cas9 RNP complexes. First, by using a dual-fluorescent reporter mT/mG mice, we show that intra-uterine electroporation of Cas9 RNP results in an efficient genomic edition of epithelial endometrial cells. Second, as a proof of its applicability for EC modeling, we show that *Pten*-targeting-RNP delivery caused loss of expression of PTEN protein in epithelial endometrial cells that resulted in the development of endometrial lesions compatible with EC. Similar results were observed by targeting other important EC driver genes such as *p53* or *Fbxw7*. Moreover, we show that the combination of CRISPR/Cas9 technology in mT/mG dual-reporter mice and light-sheet microscopy allows three-dimensional imaging and tracking of cancer cells.

In conclusion, we provide a novel methodology based on the *in vivo* somatic targeting of use of CRISPR/Cas9 that opens new possibilities for a rapid, flexible and multiplexable *in vivo* modeling of EC.

## METHODS

### Mouse models

Mice were housed in a barrier facility and pathogen-free procedures were used in all mouse rooms. Animals were kept in a 12-hour light/dark cycle at 22 °C with *ad libitum* access to food and water. All procedures performed in this study followed the National Institute of Health Guide for the Care and Use of Laboratory Animals and were compliant with the guidelines of Universitat de Lleida. Floxed homozygous PTEN (C;129S4-*Pten^tm1Hwu^*/J, hereafter called PTEN^fl/fl^); CRE:ER (B6.Cg-Tg(CAG-CRE/Esr1*5Amc/J) and reporter mT/mG (B6.129(Cg)-*Gt(ROSA)^26Sortm4(ACTB-tdTomato,- EGFP)Luo^*/J) mice were obtained from the Jackson Laboratory (Bar Harbor, ME, USA).

### Preparation of CRISPR/Cas9-RNP

CRISPR RNAs (crRNA) and trans-activating CRISPR RNA (tracrRNA) (Alt-R® CRISPR-Cas9 tracrRNA and Alt-R® CRISPR-Cas9 crRNA) were obtained from Integrated DNA Technologies (IDT, Coralville, IA). The sequences of the crRNAs used in this work are listed in the Supplementary Methods (Table S1). Lyophilized crRNAs and tracrRNA were resuspended in nuclease-free duplex buffer (100 mM potassium acetate; 30 mM HEPES, pH 7.5) (Nuclease-Free Duplex buffer, IDT) at a 100 µM concentration. To hybridize tracrRNA and crRNA were mixed at equimolar concentrations, diluted in Nuclease-Free Duplex buffer to a 20 µM concentration. Mixed RNAs were heated at 95°C for 5 minutes and subjected to a negative temperature gradient (−2 °C/30 sec) in a thermal cycler until they reach room temperature. Hybridized 20 µM stocks of crRNA:tracrRNA were stored at -20 °C.

Recombinant Streptococcus pyogenes Cas9. (Alt-R® S.p Cas9 Nuclease V3, IDT) was mixed at equimolar concentration with crRNA:tracrRNAs and diluted in Opti-MEM (Invitrogen, Waltham, MA) at a final concentration of 6 µM in 5 µL (0.5 µL of 61 µM Cas9, 1.5 µL of 20 µM crRNA:tracrRNA and 3 µL of Opti-MEM). Mixture was incubated for 20-30 minutes at room temperature to allow Cas9-crRNA:tracrRNA RNP formation (Cas9-RNP).

### *In vitro* CRISPR/Cas9 RNP assay

To test Cas9 nuclease on-target activity of *Pten*-RNP, *p53*-RNP or *Fbxw*7-RNP a PCR flanking Cas9 RNP target sequences was performed using as template genomic DNA extracted gDNA Isolation Kit (NYZTech, Lisbon, Portugal) following manufacturer’s instructions from wild-type mouse skin fibroblasts. Primers sequences and the PCR conditions for amplification of DNA fragments are specified in Supplementary Methods (Table S3). PCR amplicons were resolved in agarose gels, purified using the NZYGelpure kit (NZYTech) following manufacturers’ instructions and quantified with a NanoDrop spectrophotometer.

pCA-mT/mG vector (gift from Liqun Luo, #26123, Addgene) or *Pten*, *p53* and *Fbxw7* amplicons were submitted to cleavage assay with *LoxP*-RNP, *Pten*-RNP, *p53*-RNP or *Fbxw7*-RNP, respectively. In a 10 µl reaction volume, 2 µl containing 0.5-1 µg of purified PCR amplicons or pCA-mT/mG were mixed with the 3µl of 2µM RNP and 5 µL of Opti-MEM. Reactions were incubated for 90 minutes at 37 °C and 5 minutes at 95 °C to stop reaction. Nuclease assays were resolved in a 1% agarose gel.

### Isolation and culture of mouse skin fibroblast

Tail and ear fragments from C57/B6 or mT/mG mice were dissected and incubated in 2-3 hours at 37 °C in 950 µL of Dulbecco’s modified Eagle’s medium (DMEM, Invitrogen), supplemented with 10% inactivated fetal bovine serum (FBSi, Invitrogen), 1 mmol/L HEPES (Sigma -Aldrich, San Luis, MO), 1 mmol/L sodium pyruvate (Sigma-Aldrich), 1 % penicillin/streptomycin (Sigma-Aldrich) and 0.1 % Amphotericin B (Invitrogen) (supplemented DMEM) and 50 µL Collagenase Type IA 200 mg/mL (Worthington Biochemical Corporation Lakewood, NJ). Digested skin fragments were filtered through sterile Cell Strainer (70 µM Nylon Mesh, Fisherbrand, Hampton, NH). Cell strainer was washed with supplemented DMEM medium to recover most of digested cells. Isolated fibroblasts were centrifuged, resuspended in complete medium and plated in P100 cultures plates. Twenty-four hours after plating, medium was changed for fresh supplemented DMEM medium.

### Mouse fibroblast transfection of Cas9 RNP

mT/mG mouse fibroblast were seeded in M24 multiwell plates and at 60-70% confluency. After 24 hours, medium was replaced by 400 µl of Opti-MEM. A 0,12 µM *LoxP*-RNP mix was prepared by diluting of 6 µM *LoxP*-RNP in 25 µl of Opti-MEM. In a separate tube, 1.5 µL of Lipofectamine was added to 25 µL of Opti-MEM. Lipofectamine mix was added to RNP mix, vortexed and incubated at room temperature for 20 minutes. The 50 µl volume of the resulting mixture was added dropwise to cells. After 24 hours medium was replaced by supplemented DMEM.

### *In vivo* Cas9 RNP delivery into mouse uterus by electroporation

mT/mG mice were anesthetized with 2 % isoflurane by inhalation. A longitudinal incision in the skin and peritoneal wall in the abdomen was performed using sterile dissection scissors and forceps. Uterus was located and pulled out of the abdominal cavity. Once uterus was exposed, 5 µL of 6 µM RNP supplemented with Fast Green FCF (Sigma) to facilitate visualization of RNP complexes delivery were injected to each uterine horn using a Hamilton Neuros syringe (Hamilton, Reno, NV). For the injection of more than one Cas9-RNP complexes, 5 µl of mixture in 1:1 ratio of each Cas9-RNP was injected per uterine horn. Using a BTX830 square electroporator (BTX, Hawthorne, NY), 4 pulses of 50 mV for 50 msec spaced by 950 msec were applied to the injected uterine horn. This protocol was repeated opposing the orientation of tweezers and performed along all the entire uterine horn. The tweezers used were Platinum Tweezertrode, 5 mm Diameter (BTX). Once electroporated, uterus was carefully reintroduced back into the peritoneal cavity, and the skin and peritoneal wall was closed with a wound stapler (AutoClip System 12020-09, Fine Scientific Tools, FST, Poznań Poland). Mice were sacrificed by cervical dislocation, at different time-points post-electroporation, depending on the experiment. Uterus was dissected and processed as required for each analytical procedure described in this section.

### Isolation and sorting of endometrial epithelial cells

Isolation of mouse endometrial epithelial cells was performed as previously described (31). Mice were sacrificed by cervical dislocation, and uterine horns were dissected from electroporated mT/mG mice. Uteri were washed with Hanks (HBSS) (Invitrogen) and chopped in 3- to 4-mm-length fragments. Uterine fragments were digested with 1% trypsin (Invitrogen) in HBSS for 1 hour at 4°C and 45 minutes at room temperature. Trypsin digestion was stopped by addition of DMEM containing 10% fetal bovine serum (Invitrogen). After trypsin digestion, epithelial sheets were squeezed out of the uterine pieces by applying gentle pressure with the edge of a razor blade. Epithelial sheets were washed twice with PBS and resuspended in 1 ml of DMEM/F12 (Invitrogen) supplemented with 1 mmol/L HEPES (Sigma-Aldrich), 1% penicillin/streptomycin (Sigma-Aldrich), and Fungizone (Invitrogen) (basal medium). Epithelial sheets were mechanically disrupted in basal medium by pipetting 50 times through a 1-ml tip, until clumps of cells are observed under the microscope. Cells were diluted in basal medium containing 2% dextran-coated charcoal-stripped serum (DCC) (HyClone Laboratories, Logan, UT) and plated into culture dishes (BD Falcon, Bedford, MA). Cells were cultured for 24 hours in an incubator at 37°C with saturating humidity and 5% CO_2_.

### NGS-amplicon sequencing analysis

Electroporated uteri were dissected and opened longitudinally to expose the endometrial epithelial cells of the uterine lumen. Under a fluorescence stereoscopic microscope (Nikon Eclipse Ts2R), luminal epithelial areas displaying green cells were rinsed with HBSS (Invitrogen) and scraped with a scalpel blade. Scraped sheets of epithelial cells were collected rinsed with PBS based on a series of centrifugations. Cell pellet was processed to extract the genomic DNA from the endometrial epithelial cells using the NZY Tissue gDNA Isolation Kit (NYZTech), following the manufacturers’ instructions. Genomic DNA was amplified by PCR using primers flanking RNP-targeted region. Primer sequences and PCR conditions for amplification of DNA fragments are specified in Supplementary Methods (Table S3). PCR amplicons were resolved in agarose gels, purified using the NZYGelpure kit (NZYTech) following manufacturers’ instructions and quantified with a NanoDrop spectrophotometer. Next Generation Sequencing (NGS)-Amplicon sequencing was carried out by Genewiz (Azenta Life Sicences, Chelmsford, MA) company. Raw FASQ archives containing NGS-amplicon sequences were submitted to bioinformatic analysis to detect the presence of indels using Crispresso2 (32) and Cas-Analyzer (33) software.

### Tissue processing and immunohistochemistry analysis on paraffin sections

Animals were euthanized and uteri were collected, formalin-fixed O/N at 4 °C and embedded in paraffin for histologic examination. Three µm sections of the paraffin blocks were dried for one hour at 80 °C, dewaxed in xylene, rehydrated through a graded ethanol series, and washed with phosphate-buffered saline. Antigen retrieval was performed in EnVision™ FLEX, High pH or Low pH solution (DAKO, Glostrup, Denmark) for 20 minutes at 95°C, depending on the primary antibody. Endogenous peroxidase was blocked by incubating slides with 3% H2O2 solution. After three washes with phosphate-buffered saline (PBS) (DAKO), primary antibody was incubated for 30 minutes at room temperature, washed in PBS and incubated with secondary antibody. A peroxidase (HRP)-conjugated secondary antibody or a biotin conjugated secondary antibody plus peroxidase-linked streptavidin were used depending on the primary antibody used. The reaction was visualized with the EnVision Detection Kit (DAKO) using diaminobenzidine chromogen as substrate. Sections were counterstained with Harris haematoxylin. Antibodies used in this study and immunohistochemistry conditions are summarized in Supplementary Methods (Table S4)

The samples were stained with Hematoxylin and Eosin (H-E), reviewed and evaluated histologically by two pathologists, following pre-established uniform criteria. Representative images were taken with Leica DMD 108 microscope.

### Uteri optical clearing

For three-dimensional imaging of mT/mG mice, uteri from mice electroporated with *LoxP*-RNP with and without *Pten*-RNP were dissected and lightly fixed in 4% paraformaldehyde (PFA) in PBS (pH 7-7.4) for 2-3 hours at 4 °C, to preserve the structure of the tissue without losing fluorescence. After washing with PBS for 30 minutes, uteri were cleared with a variant of the CUBIC chemical clearing protocol (34). In order to preserve structural conformation and fluorescence of mGFP, tissue rinsing and clearing steps were carried out under aqueous conditions as previously described (34). Tissues were directly immersed in CUBIC-L solution (10 wt % N-Butyldiethanolamine [Tokyo Chemical Industry, B0725], 10 wt % Triton X-100 [Merck, 1.08603.1000] and 80 wt % MilliQ water). Uteri samples in CUBIC-L were incubated with gentle shaking at 37 °C for 7 days in total. After 4 days of incubation, the CUBIC-L was refreshed once. After delipidation, tissues were washed in PBS with shaking at room temperature for 6 hours at least. Throughout all the protocol, samples were preserved from light exposure to avoid fluorescence loss.

### Lightsheet Fluorescence Microscopy

After washing with PBS for 30 minutes, uteri samples were embedded into a 0.8% solution of low-melting agarose (A9045, Sigma) diluted in MilliQ water to enable easy mounting of the sample in our custom lightsheet system. For refractive index (RI) matching between the sample and the embedding agarose, the blocks containing the tissue were incubated at room temperature overnight in 1:1 water-diluted CUBIC-RA (a mixture of 45 wt% antipyrine [Tokyo Chemical Industry, D1876], 30 wt% N-methylnicotinamide [Tokyo Chemical Industry, M0374)] and 25 wt% MilliQ water). Tissues were then immersed in CUBIC-RA with gentle shaking at room temperature for 2-3 days before imaging.

Once RI was matched, three-dimensional visualization of the entire tissue was carried out using lightsheet fluorescence microscopy. For the mesoscopic analysis of the uteri electroporated with RNP CRISPR/Cas9, lightsheet fluorescence macroscopy based on a custom instrument (MacroSPIM, see details in Kennel et al., 2018)(35) was used. In brief, the agarose block containing the tissue was placed in a quartz glass cuvette and immersed in the same CUBIC-R used to match the RI between agarose and tissue. Imaging was performed at a magnifications of 7.2x (Figure 3) and 4.8x (Figure 6), yielding respective pixel size XY = 0.902 and 1.345 µm, and with Z steps of 2.5 and 5µm resepectively. The lightsheet waist was adjusted to be around 4.5 to 5 µm for axial resolution. Fluorescence of tdTomato was excited with a 561 nm laser, and collected with a bandpass filter BP609/54, and fluorescence of GFP was induced with a 488 nm line, collected with a BP525/50 (Semrock filters).

### Image processing, analysis and visualization

Raw images were first processed with the open-source MosaicExplorerJ (36) macro running in Fiji (37) for the stitching of large mosaic/tiles. Image processing was done with Fiji: line plot in Figure 3 was performed with a 10pixel-wide line. Segmentation of GFP+ lesions was performed with a custom ImageJ1 macro that, in brief, enables to i) generate regions of interest (ROIs) using a combination of Magic Wand and Lasso tool every 10 planes, ii) interpolate between those ROIs, and iii) save the masks in 16bit tiff format for further visualization. Three-dimensional visualization was performed with the free Imaris Viewer (Bitplane, Oxford Instrument), including surface rendering in “Normal Shading” mode in Figure 3J and 6F. Imaris 9.1 (Bitplane, Oxford Instrument) was used only to generate smoothed surfaces of the lesions masks (Figure 6F,K) and supplementary videos.

## RESULTS

### Design of a reporter method to monitor CRISPR/Cas9 edition in living cells

To follow-up CRISPR/Cas9 activity in living cells, we first designed an easy method for identification of the edited cells. First, we tested our approach *in vitro* using as a target DNA, a plasmid that contains a *LoxP*-flanked membrane-targeted tandem dimer Tomato (mT) followed by membrane-targeted GFP (mG) under the control of constitutive promoter (pCA-mT/mG, Figure A). Upon Cre-directed recombination, *LoxP*-flanked mT is deleted and recombined cells shift from red to green fluorescence. Therefore, we designed a crRNA targeting a common PAM sequence adjacent to both *LoxP* sites flanking mT (Figure 1A). Next, we incubated the pCA-mT/mG plasmid with the RNP formed by the crNA:tracrRNA targeting *LoxP* plus recombinant Cas9 and we check the Cas9 mediated cleavage of the target DNA in an agarose gel (Figure 1A). As expected, we observed two bands, a 2.45 Kb DNA fragment containing the mT-pA sequence and a 6 kDa DNA fragment corresponding to the rest of the plasmid, indicating successful cleavage of Cas9 at the expected sites (Figure 1B). Noteworthy, the effect of the RNP was specific, as no cleavage was observed when only one of the two components, the guide RNA or the Cas9, was present (Figure 1B).

**FIGURE 1.**
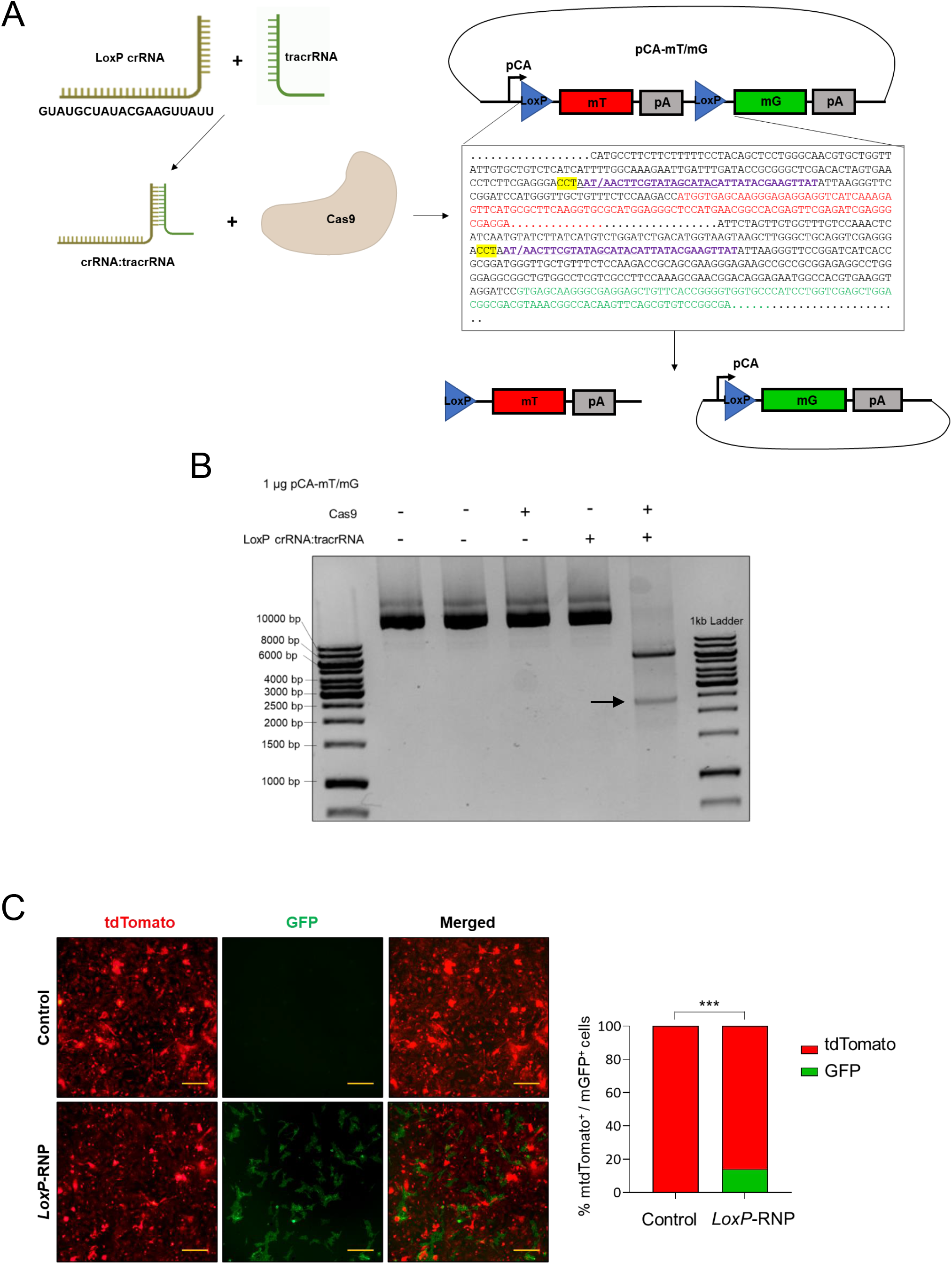
Design of *LoxP*-targeting RNP to monitor CRIRP/Cas9 activity. **(A)** Representative diagram CRISPR/Cas9 mediated edition of *LoxP* sites on the pCA-mT/mG reporter plasmid. PAM sequences highlighted in yellow, the crRNA sequence targeting *LoxP* site underlined in purple, *LoxP* sequences in blue, part of the coding sequence for the fluorescent protein tdTomato (mT) is in red, part of the coding sequence for the fluorescent protein EGFP (mG) in green. pCA; promoter formed by Cytomegalovirus enhancer and chicken β-Actin promoter. pA; SV40 polyadenylation signal. **(B)** Representative image of an agarose-resolved *in vitro* CRISPR/Cas9 digestion of pCA-mT/mG. Arrowhead indicates the 2.45 Kb band corresponding to *LoxP*-flanked mT. **(C)** Representative images and quantification of red and green cells of mT/mG^f/f^ mouse fibroblast cultures 48h after being transfected with *LoxP*-targeting RNP (*LoxP*-RNP). Images at 10X magnification. Scale bars 100µm.

Next, we tested the ability of our *LoxP*-targeting Cas9 RNP to induce gene edition into living cells. For this purpose, skin fibroblasts isolated from mT/mG reporter mouse (38) were transfected with the *LoxP*-targeting Cas9 RNP, and the number of green (edited) or red (non-edited) cells was quantified 48 hours after transfection (Figure 1C). As observed, *LoxP*-targeting Cas9 RNP caused a switch from red-to-green in ∼10% of total fibroblasts, indicating a successful edition in living cells in culture.

### CRISPR/Cas9 electroporation *in vivo* causes genomic edition of mouse endometrial epithelial cells

The above results enabled us to test the ability of our *LoxP*-targeting Cas9 RNP *in vivo*, in live animals. Again, we took advantage of the mT/mG reporter mouse (38) and try to target endometrial epithelial cells by intrauterine transfection of our *LoxP*-targeting Cas9 RNP. Briefly, abdominal cavity of anesthetized mT/mG females was opened longitudinally, and the uterine horns were exposed. Then, 5 µl of *LoxP*-targeting Cas9 RNP mix containing 0,5 µl of 62 mM recombinant Cas9, 1,5 µl of 20 µM crRNA:tracrRNA duplex and 2µl of Opti-MEM and 1 µl of Fast Green were loaded into a Hamilton Neuros Syringe and injected into one uterine horn. Transfection of the RNP was achieved by electroporation using a BTX830 square electroporator and applying 4 pulses of 50 mV during 50 msec spaced by 950 msec gap to the injected uterine horn (Figure 2A). This protocol was repeated in the same region of the uterus opposing the orientation of the tweezers and repeated along all the uterine horn. After 15 days, uterine horns were dissected, opened longitudinally and examined for the presence of green cells under a fluorescent stereoscope (Figure 2B). Interestingly, only uterine horns electroporated with *LoxP*-targetingCas9 RNP displayed cells with GFP expression, indicating that genome edition was achieved into the uterine cavity (Figure 2B). As controls, Cas9 alone or crRNA:tracrRNA alone were injected and electroporated or *LoxP*-targeting RNP was injected without administration of electric pulse. None of these conditions showed the presence green cells 15 days after the protocol. To further demonstrate the correct genomic edition of *LoxP* sites *in vivo*, endometrial cells of the uterine horn displaying green cells were carefully dissected, submitted to genomic DNA extraction, and amplification by PCR with primers flanking *LoxP* sequences. Cas9 cleavage of both LoxP-targeted sites and end joining would give a 181 bp band, in which insertions or deletions caused by NHEJ could be observed (Figure 2C). Indeed, PCR amplification rendered an abundant product of 181 bp compatible with both *LoxP* excision and subsequent joining (Figure 2C). To check for presence of indels in joined sequences, PCR products were submitted to NGS-amplicon sequencing. NGS results were analyzed with the Cas9 edition detection software Crispresso2 (32) and Cas-Analyzer (33) with similar results. Using as a reference the predicted genomic sequence in which a perfect joining between the two Cas9 generated ends after mT excision (*LoxP*-edited mT/mG, Figure 2C), we found that most sequences (90.92%) aligned with the reference sequence, indicating that Cas9-generated ends healed without indels. Only a small percentage of sequences (9.08%) displayed different types of indels (Figure 2C).

**FIGURE 2.**
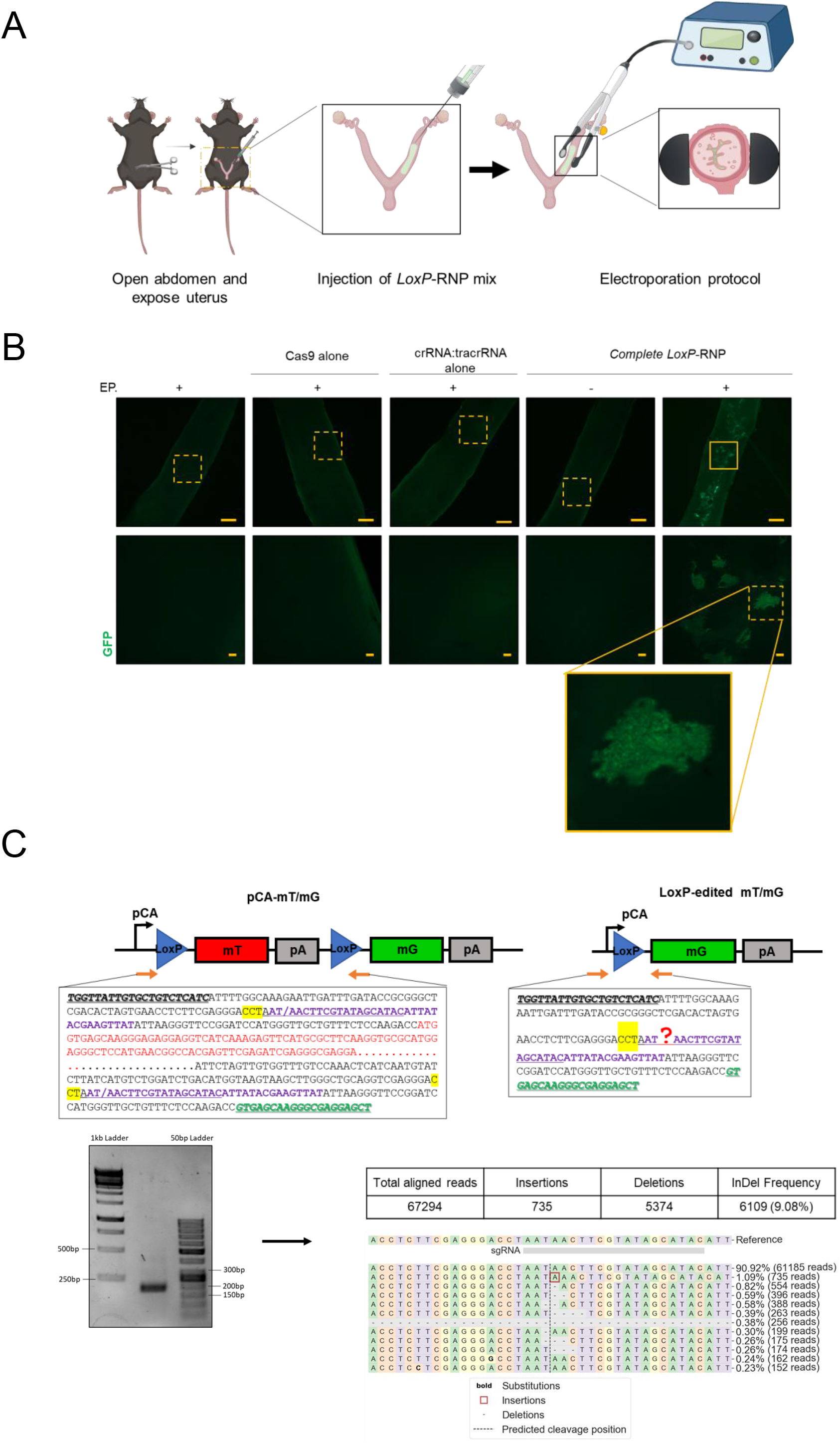
Protocol for Cas9 RNP delivery by electroporation of into mouse uterus. **(A)** Diagram depicting mouse surgery, intra-uterine RNP delivery and electroporation protocol. **(B)** Fifteen days after electroporation in the uterine cavity of mT/mG^f/f^ mice, the green fluorescence of the endometria, indicative of nuclease activity, was analyzed. Representative images of the uterine branch where gene editing is only seen in the condition where complete *LoxP*-RNP was injected, and the electric pulse was applied (EP). In experimental conditions in which Cas9 or crRNA:tracrRNA alone were injected or in the condition in which complete *LoxP*-RNP was injected but no EP was applied, no green cells observed. n=4 uterine horns per condition. Images at 2X and 10X magnification. Scale Bars: 1mm and 100um, respectively. **(C)** Representative diagram of predicted changes of the mT/mG reporter mouse construct before and after CRISPR/Cas9 mediated edition of *LoxP* sites.PAM sequences highlighted in yellow, the crRNA sequence targeting *LoxP* site underlined in purple, *LoxP* sequences in blue, part of the coding sequence for the fluorescent protein tdTomato (mT) is in red, part of the coding sequence for the fluorescent protein EGFP (mG) in green. pCA; promoter formed by Cytomegalovirus enhancer and chicken β Actin promoter. Primers for are in bold. Representative agarose gel showing 181bp amplicon corresponding to DNA fragment flanking *LoxP* sites. Summary of Cas-analyzer software analysis of NGS-amplicon sequencing of purified agarose band. Upper table indicate the number of unmodified reads and reads displaying insertions or deletion (indels). Bottom table indicate the 12 most frequent indels compared to reference sequence. Horizontal dashed lines indicate deleted sequences. The vertical dashed line indicates the predicted cleavage site.

### *In vivo* CRISPR/Cas9 genomic edition is restricted to endometrial epithelial cells

Endometrium contains epithelial endometrial and stromal cells surrounded by the myometrium. Therefore, we analyzed which cell types were edited by the electroporated *LoxP*-targeting Cas9 RNP. To address this point, we performed immunohistochemistry with antibodies against cytokeratin-8 (CK6) and GFP on consecutive paraffin sections of the electroporated uterine horns. In all paraffin sections analyzed, expression of GFP was restricted to cytokeratin expressing cells, indicating that genomic edition was restricted to endometrial epithelial cells (Figure 3A-B). To further demonstrate an epithelial-specific genomic edition, we performed a whole uterine analysis of GFP expression using light sheet microscopy. First, we performed an image-based analysis of GFP expression using lightsheet microscopy on an optically cleared whole uterine fragment electroporated with *LoxP*-targeting Cas9 RNP. Optically sectioned images and three-dimensional reconstruction of an approximately 7mm long uterine tissue revealed that GFP edition was restricted to cells displaying epithelial sheet of glandular organization (Figure 3C-H, Supplementary video 1). Lighsheet imaging revealed the tissue-wide mosaic pattern distribution of the GFP expressing cells, and confirmed the expected deletion of *LoxP*-flanked mT (tdTomato) in those cells, as shown in Figure 3K-M.

**FIGURE 3.**
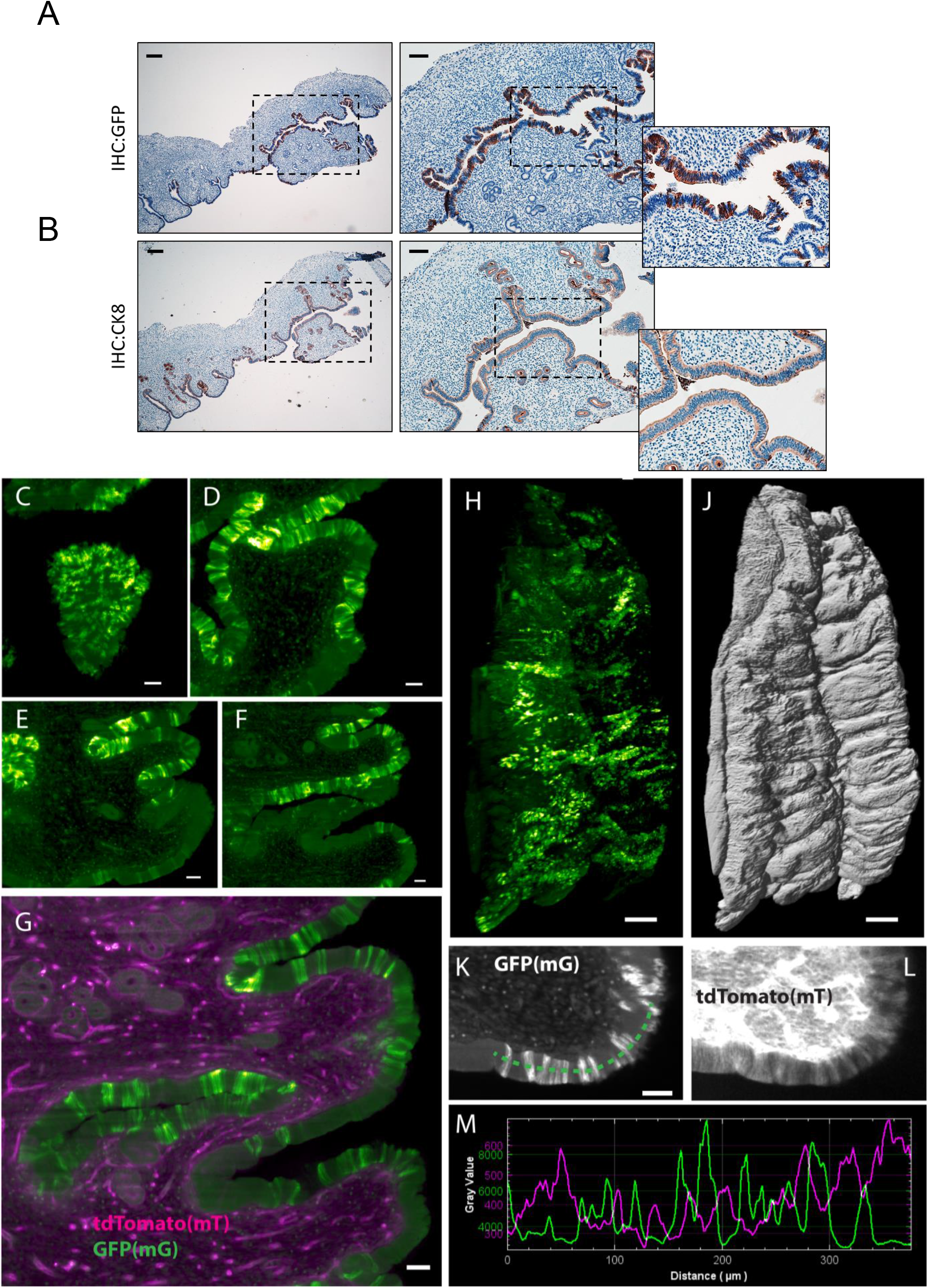
Cas9 RNP electroporation of mouse uterus leads to specific edition in endometrial epithelial cells. Representative images cytokeratin 8 (CK8) **(A)**, and GFP **(B)** of consecutive histological sections of mT/mG^f/f^ uteri 15 days after electroporation of *LoxP*-RNP **(C-L)** Representative light sheet microscopy images corresponding to mT/mG^f/f^ endometria 15 days after electroporation with *LoxP*-RNP. (**J**) shows the GFP cells distribution in a 7mm-long endometrium, **(H)** the surface of the endometrium detected from the tdTomato channel. **(C to G)** are subsequent close-up sections in a representative XY location at Z depths, calculated from the tissue surface, of 20µm (C), 108µm (D), 232µm (E), 375µm (F), 450µm (G), where a GFP+ mosaic pattern can be seen in the epithelial cells of the endometrium. **(G)** shows the merged fluorescence acquired with the green GFP channel with overlapped tdTomato shown in magenta. K-L show the respective distribution of GFP and tdTomato cells along the endometrium epithelium. **(M)** shows the complementary intensity profile (see peaks to valley correspondence), measured on K (green) and L (magenta), along the green dashed line displayed in K). Images (A-B) were acquired at 4X and 10X magnification. Scale bars: (A), 200µm; (B), 100µm; (C-G), (K-L), 50µm, (H-J), 500µm. (H-L) was reconstructed from a vertical tiled image 1×5 fields, all images were acquired at a magnification of 7.2x with a lightsheet thickness of approximately 4.5µm (C-G) are representative capture fields corresponding to the Supplementary video 1.

### *In vivo* electroporation of Cas9 RNP causes genomic edition of frequently mutated driver genes in EC

The above results enabled us to design a genomic edition protocol for targeting genes involved in EC. According to the whole genome sequencing of EC (27) we selected three of the most frequently mutated genes: *Pten*, *p53* and *Fbwx7*. Then we designed two different crRNAs targeting exon 5 of *Pten* and one crRNA targeting either *p53* or *Fbxw7* (sequences are provided in supplementary methods). To test the nuclease activity of *Pten*, *p53* and *Fbxw7* Cas9 RNP complexes, we first performed an *in vitro* Cas9 cleavage assay (Figure 4A). In this case, as a target DNA we used PCR amplicons of the genomic region targeted by our crRNAs (Figure 4A). All Cas9 RNPs were able to cleave the corresponding PCR rendering the expected size fragments (Figure 4A). Because PTEN2 crRNA produced sharper and cleaner cleavage bands, we selected this crRNA for further experiments (from now on, we will refer as *Pten*-targeting-RNP or *Pten*-RNP the one containing crRNA2).

**FIGURE 4.**
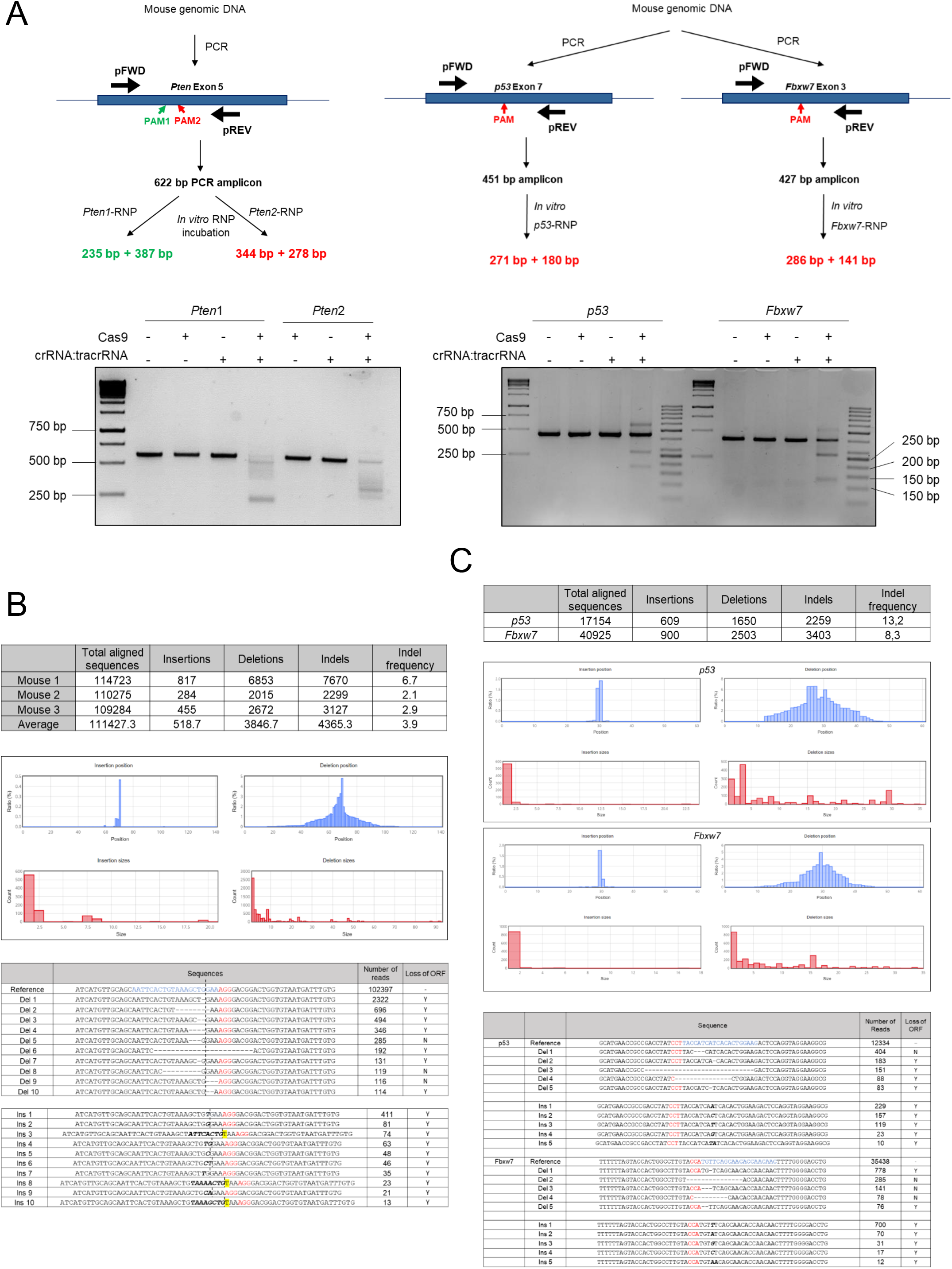
*In vivo* CRIRP/Cas9 mediated gene edition of *Pten*, *p53* and *Fbwx7* in mouse endometrial cells. **(A)** Representative diagrams and agarose gels of *in vitro* CRISPR/Cas9 activity assay and to evaluate *Pten*, *p53* and *Fbxw7* targeting RNPs. **(B)** Bioinformatic analysis of amplicon sequences from *Pten*-RNP, *p53* and *Fbxw7* electroporated mice. Analysis of NGS-*Pten* amplicons with Cas-analyzer software. Upper table indicate the number of unmodified reads and reads displaying insertions or deletion (indels) in three different mice. Graphs indicated the position and size of indels in a region spanning 70 nt each site of predicted cutting site. Bottom table indicate the top ten insertions and deletions compared to reference sequence, including the number of read and whether they cause (Y) o not (N) loss of Pten open reading frame. PAM are in red, crRNA sequence in blue. Horizontal dashed lines indicate deleted sequences. The vertical dashed line indicates the predicted cleavage site. Bold/cursive characters indicate insertions. Yellow highlighted indicate substitutions nearby insertion sites. **(C)** Analysis of NGS-*p53* and *Fbxw7* amplicons with Cas-analyzer software. Upper table indicate the number of unmodified reads and reads displaying insertions or deletion (indels) in three different mice. Graphs indicated the position and size of indels in a region spanning 70 nt each site of predicted cutting site. Bottom table indicate the top five insertions and deletions compared to reference sequence, including the number of read and whether they cause (Y) o not (N) loss of *p53* and *Fbxw7* open reading frame. PAM are in red, crRNA sequence in blue. Horizontal dashed lines indicate deleted sequences. The vertical dashed line indicates the predicted cleavage site. Bold/cursive characters indicate insertions.

Next, we evaluated the ability of gene editing of these RNPs to cause genomic edition of endometrial cells *in vivo*. For this purpose, females were electroporated with *Pten*, *p53* or *Fbwx7* RNPs and sacrificed 15 days after electroporation. Endometrial cells were dissected from the uterus and sequences flanking crRNA targets were amplified by PCR and submitted to NGS-amplicon sequencing. NGS analysis of *Pten* exon 5 amplicons from three different mice demonstrated the presence of insertion or deletions (indels) surrounding predicted RNP cleavage site (Figure 4B). Similarly, NGS-amplicon sequencing analysis of mice electroporated *p53* and *Fbwx7* RNPs demonstrated the presence of gene edition in these loci (Figure 4C). Percentages of edition were variable due to two different determinants: the efficiency of each RNP complex to induce target cleavage and the electroporation efficiency in each individual mouse.

### *In vivo* CRISPR/Cas9-mediated *Pten* knock-out on endometrial epithelial cells leads to development of endometrial neoplasia

Among all mutated genes, *Pten* is the tumor suppressor gene displaying the highest frequency of mutation in EC (25, 26). In mice, the role of *Pten* in EC has been extensively demonstrated by many *Pten* conditional knock-out mouse models using the *LoxP* flanked exon 5 allele (39)(21). For instance, we have previously demonstrated that crossing this conditional knock-out mouse with a tamoxifen inducible Cre:ER^T^ mouse leads to loss of *Pten* expression in endometrial epithelial cells resulting in the development of EC (40). Considering these facts, we wanted to set a proof of principle for our CRISPR/Cas9 *in vivo* approach as a useful tool to mimic frequent EC mutations and recapitulate their phenotypic effects. Therefore *Pten*-targeting Cas9 RNP was electroporated in B6/B57 wild type mice. First, we tested whether CRISPR/Cas9-mediated edition of *Pten* lead to the alteration of its open reading frame that results in loss PTEN protein expression. Thus, 15 days after electroporation with *Pten*-targeting Cas9 RNPs, mice were sacrificed and PTEN expression was analyzed by immunohistochemistry with specific antibodies (Figure 5A). PTEN staining revealed a mosaicism of epithelial cells showing loss of PTEN among others retaining PTEN expression. This result suggests that cells in which *Pten*-RNP was delivered and caused *Pten* InDels compatible with loss of its open reading frame, lose PTEN protein expression. Next, we analyzed whether CRISPR/Cas9-induced loss of PTEN expression was enough to initiate endometrial carcinogenesis. For this purpose, mice were sacrificed 12 weeks after electroporation, and endometrial fragments of uterine horns were dissected and submitted to histopathological evaluation. Interestingly, hematoxylin-eosin staining revealed areas of endometrial epithelial neoplasia (CK8 positive but with no PTEN expression) surrounded by normal epithelium (CK8 and PTEN positive) (Figure 5B). suggesting that CRISPR/Cas9 genomic edition of *Pten* in individual epithelial cells may cause an increase of PI3K/AKT signaling pathway activity which has been reported to be a key step during development of EC. To address this hypothesis, we performed an immunohistochemistry analysis of serial endometrial sections with cytokeratin 8 (CK8) antibody to determine epithelial nature of edited cells and with PTEN and p-AKT antibodies to reveal cells in which edited Pten locus resulted in loss of PTEN protein expression and subsequent activation of AKT (Figure 5B). These results demonstrate that intrauterine *in vivo* CRISPR/Cas9 genomic edition of *Pten* in individual epithelial cells recapitulates the signaling events that lead to EC and that can be used as a novel approach to study EC pathogenesis.

**FIGURE 5.**
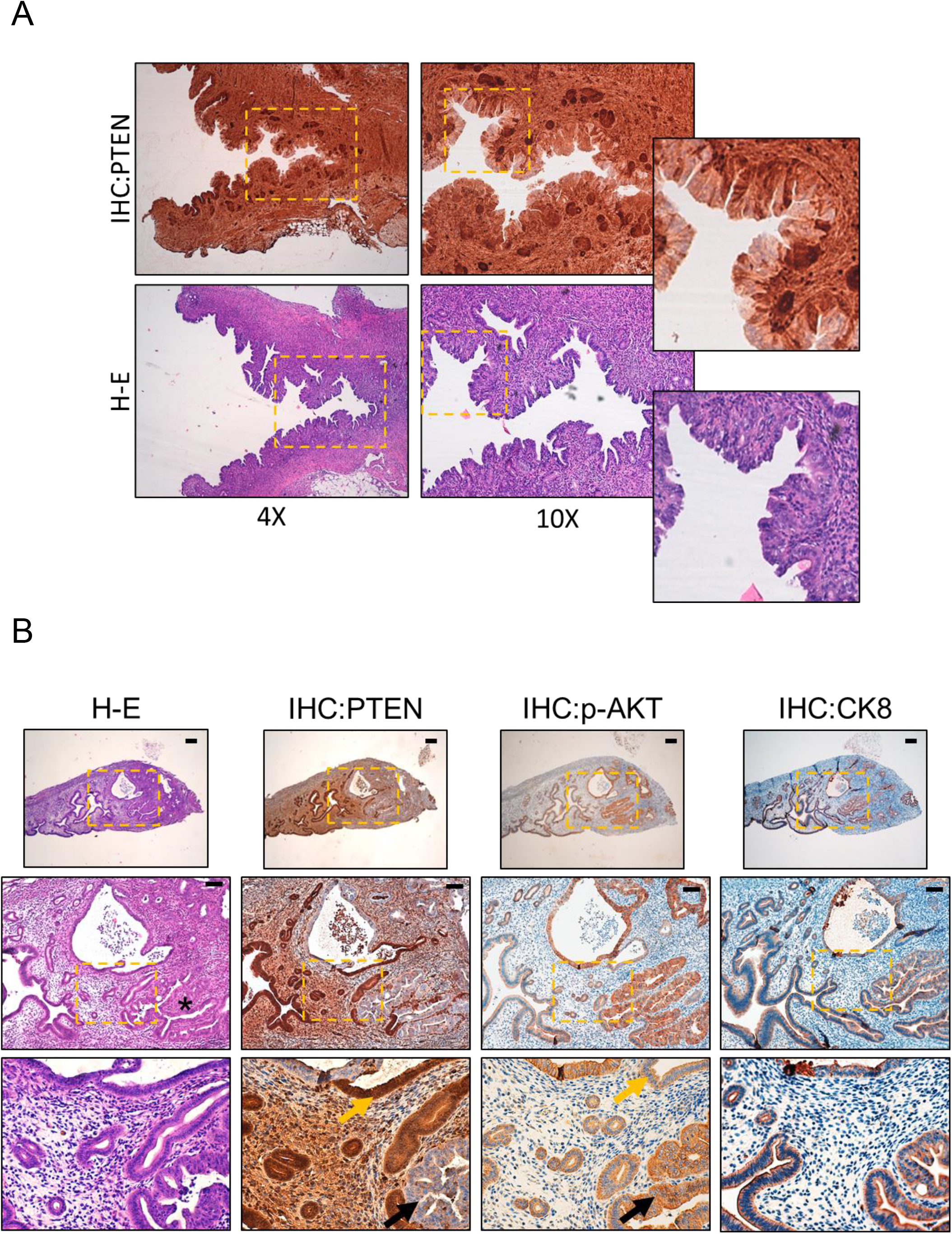
*In vivo* electroporation of *Pten*-RNP results in efficient edition of *Pten* locus and development of endometrial lesions. **(A)** Representative images of hematoxylin-eosin (H-E) staining, and immunostaining to detect PTEN on serial histological sections of C57/B6 mice 15 days after electroporation with Pten-RNP **(B)** Representative images of hematoxylin-eosin (H-E) staining, and immunostaining to detect PTEN, phosphorylated AKT (p-AKT), cytokeratin 8 (CK8), on consecutive histological sections of C57/B6 mice uteri 12 weeks post-electroporation with *Pten*-RNP. Yellow arrows indicate PTEN expressing wild type epithelial cells edited in *LoxP* locus (GFP positive). Black arrows indicate tumoral cells lacking PTEN expression that have been edited in *LoxP* locus (GFP positive). Yellow asterisks indicate tumoral cells lacking PTEN expression that have without edition in *LoxP* locus (GFP negative). Black asterisks indicate PTEN expressing wild type cells without edition in *LoxP* locus (GFP negative).

### *In vivo* CRISPR/Cas9-mediated *LoxP* and *Pten* edition generates heterogenous endometrial cell populations

One of the main advantages of the Cas9 system is its ability to simultaneous edit more than one locus at a time (4, 6). By co-electroporating two or more RNPs targeting different loci at once, it is feasible to create heterogeneous populations of cells harboring different combinations of edited genes, which may be a valuable tool to model tumoral heterogeneity in EC. To test this hypothesis, *LoxP*-targeting and *Pten*-targeting RNPs were co-electroporated in mT/mG mice and 12 weeks later, the uteri were submitted to an immunohistochemistry (IHC) analysis on serial endometrial sections with anti-GFP antibodies to detect cells edited for *LoxP* sites, PTEN and p-AKT antibodies to reveal cells edited in the *Pten* locus and CK8 antibody to determine epithelial nature of edited cells (Figure 6A-E). Interestingly, IHC revealed heterogenous staining compatible with four different scenarios. A first group of cells that only received *LoxP*-targeting RNP, resulting in normal epithelial cells displaying positive staining for GFP and PTEN but negative for p-AKT (CK8^+^;GFP^+^;PTEN^+^;p-AKT^-^). A second group of cells that only received *Pten*-RNP, resulting in tumoral epithelial cells displaying negative staining for GFP and PTEN but positive for p-AKT (CK8^+^;GFP^+^;PTEN^-^;p-AKT^+^). A third group of cells which received both *LoxP*- and *Pten*-targeting RNPs, resulting in tumoral epithelial cells displaying positive staining for GFP and p-AKT but negative staining for PTEN (CK8^+^;GFP^-^;PTEN^+^;p-AKT^-^). Finally, a fourth group of epithelial cells that did not receive any on the two targeting Cas9 RNPs displayed negative for GFP and p-AKT staining and positive staining for PTEN (CK8^+^;GFP^-^; PTEN^+^;p-AKT^-^). It is worth to mention that in any case we did not find any tumoral cell positive for PTEN and negative for p-AKT. These results demonstrate that co-electroporation of different targeting Cas9-RNPs against different loci generate heterogenous populations of edited cells.

**FIGURE 6.**
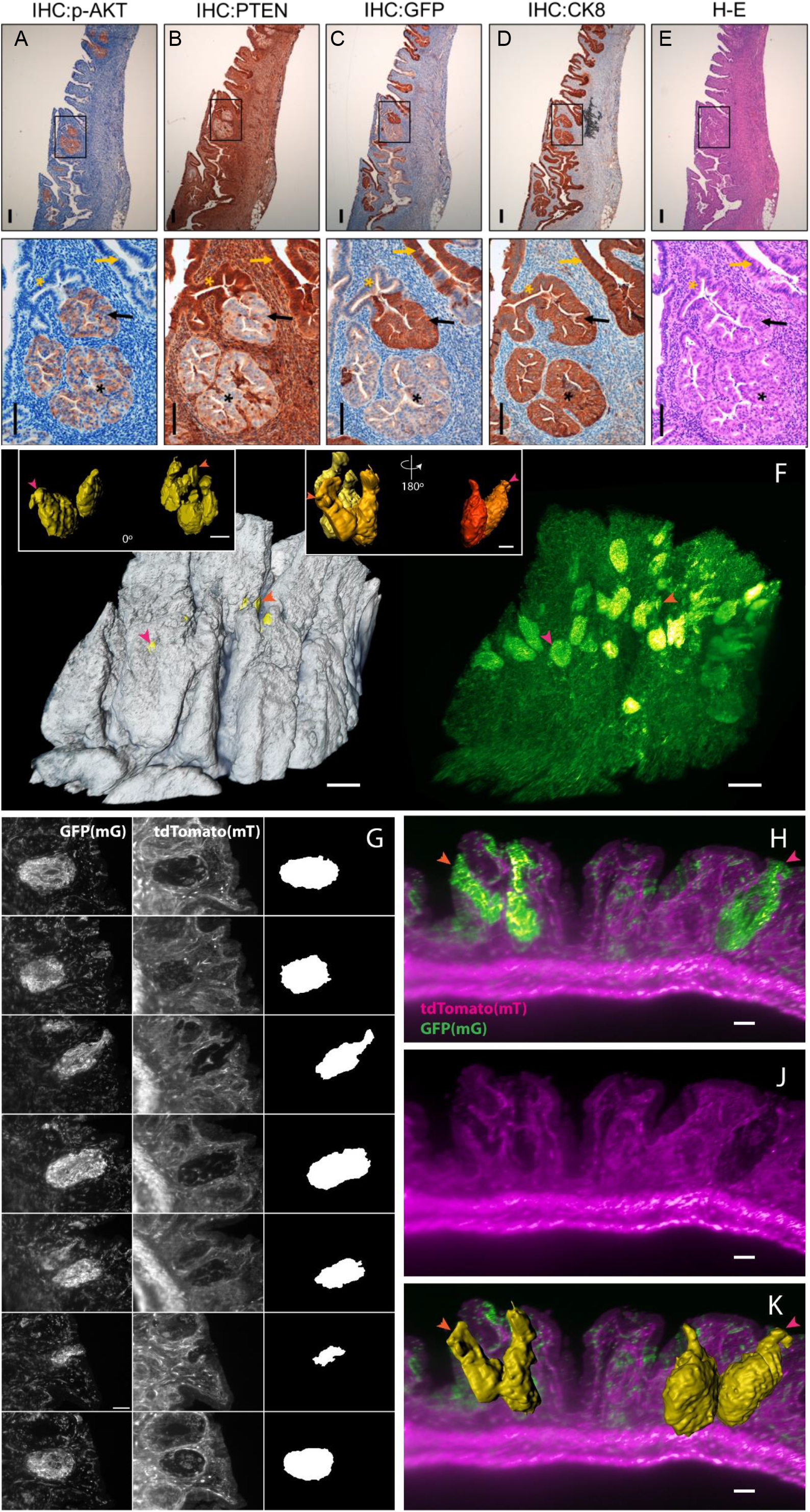
*In vivo* co-electroporation of *LoxP*-RNP and *Pten*-RNP leads to heterogenous endometrial epithelial populations. **(A-E)** Representative images of hematoxylin-eosin (H-E) staining (C), and immunostaining to detect PTEN (E), phosphorylated AKT (p-AKT) (D), cytokeratin 8 (CK8) (B), and GFP (A) of consecutive histological sections of mT/mG uteri 12 weeks post-electroporation of Pten-RNP and *LoxP*-RNP. Images show the co-existence of PTEN-deficient endometrial carcinomas either positive (black arrow) or negative (black asterisk) for GFP expression with PTEN expressing non-tumoral epithelial cells either positive (yellow arrow) or negative (yellow asterisk) for GFP expression. Images (A-E) at 4X and 20X, with scale bars 200 µm and 100 µm bars, respectively. **(F-K)** Light sheet fluorescence microscopy shows that *in vivo* CRISPR/Cas9 electroporation of *Pten*-RNP and *LoxP*-RNP uteri generates endometrial epithelial lesions throughout the tissue. Representative images and reconstructions of mT/mG endometria 12 weeks after electroporation with *Pten*-RNP and *LoxP*-RNP. **(F)** surface rendering (left, from tdTomato channel) and corresponding maximum intensity projection (right, from GFP channel) in a 3×3mm tissue fragment. Insets show six selected segmented GFP+ lesions by surface rendering, from two opposite directions: front (left, yellow surfaces) and back view (right, volumes with different colors). **(G)** shows selected sections of GFP+ lesions (left) and corresponding tdTomato signal (center), with the segmented mask of the lesions. **(H-L)** Show an orthogonal cut through three lesions, visualizing tdTomato (J), overlapped tdTomato/GFP channels, and volume rendering of the lesions (K). **(F)** and **(H-L)** clearly show the pear-shaped lesions with the tip originating from the epithelium surface, in particular see arrows in F and H. Magenta and orange arrows point at the same lesions across images F, H and K. Scales: F, 400µm, insets: left 200µm, right 100µm, G-L, 100µm. Three-dimensional video of the same sample is shown in Supplementary video 2.

Finally, we performed a lightsheet microscopy based analysis of mGFP expression on a whole uterine fragment electroporated with *LoxP*- and *Pten*-targeting RNPs. As mentioned above, some cells are co-edited with *LoxP* and *Pten* RNPs, leading to generation of GFP positive/PTEN negative cells that develop endometrial neoplasia. The observation of GFP expressing cells may be an interesting approach to monitor and trace the development of EC in CRISPR/Cas9 edited uterine cavity. Twelve weeks after electroporation with *LoxP-* and *Pten*-RNPs, endometrial tissues were fixed, cleared and imaged under lightsheet microscopy. Individual optical section images and three-dimensional reconstructions of uterine tissue revealed the growth of GFP expressing neoplasia (Figure 6F-L, Supplementary video 2), and in particular the 3D “pear” shape of those lesions originating from the endometrium surface, as shown by GFP+ segmentation and 3D rendering (Figure 6F, K).

## DISCUSSION

To date, there is no existing method for *in vivo* genomic engineering of endometrial epithelial cells. Here, we describe an easy, rapid, flexible, multiplexable and robust method for EC modelling using *in vivo* delivery of CRISPR/Cas9 by electroporation in the mouse uterus. To validate this method for EC modelling, we targeted *Pten*, the main tumor suppressor gene involved in EC.

In the present study, we have used Cas9 RNPs electroporation as a method for *in vivo* delivery of Cas9 and gRNA into the mouse uterine cavity. Delivery of targeting Cas9 RNP complexes offers some advantages over the use of plasmids encoding Cas9 and guide gRNAs. First, RNPs can immediately edit target genes without the need of transcription of translation, allowing a rapid genomic edition upon its intranuclear delivery. Second, delivery of Cas9 RNP complexes is rapid and transient, rendering higher edition rates and reduced off-targets, insertional mutagenesis and toxicity (41–43). The use of DNA-based strategies to deliver Cas9 machinery into the cells by either transient transfection or using viruses enhance the risk of accidental genomic integration leading to insertional mutagenesis (44, 45). Third, DNA-based delivery of Cas9 and gRNA can lead to prolonged and uncontrolled expression of Cas9 and guide RNAs, increasing the chances of off-target effects, and disrupting endogenous genes (46, 47). Finally, introduction of exogenous DNA sequences can boost host immune response, limiting its use for therapeutic applications (48). Despite the advantages of Cas9 RNP, its delivery *in vivo* is still a challenging issue. Direct transfection of RNP has been achieved using a wide variety of delivery systems such as microinjection, lipid-based transfection, nanoparticles, electroporation and a long etcetera of emerging materials and techniques (18,47,49). Among physical methods, *in vitro* electroporation has been widely used for RNP transfection in cell lines, primary cultures or explants (49). Although microinjection and electroporation of Cas9 RNP in to mouse embryos has been used to efficiently generate knock-out mice (50–53), *in vivo* electroporation of Cas9 RNPs to generate adult somatic GEMM has never been used. Instead, systemic delivery of RNP using nanoparticles has been recently used to edit tissues including muscle, brain, liver, and lungs (54). Lately, lipid based strategies (55, 56) or different types of nanoparticles and vesicles such as nanoclews (57), carbolylated nanoparticles (58), gold nanoparticles (59), nanosheets (60), extracellular vesicles (61), or VSV-typed vesicles (62), has been efficiently used for *in vivo* genome modification However, *in vivo* accurate delivery of RNP into the mouse uterus has never been achieved by any of the methods described. Our electroporation protocol allows a cell specific and a spatial-temporal controllable method to introduce mutations into endometrial epithelial cells, which is of pivotal importance for precise EC modeling. By injecting RNP complexes into the uterine luminal cavity, the luminal epithelial sheet lining on the mouse endometrium is exposed to RNP complexes and which facilitates their transfection by electroporation. It is worth to mention that EC arise from the epithelial compartment.

Another advantage of Cas9 system is its multiplexable genome engineering capacity that allows simultaneous edition of more than one locus at a time. Since the ability of the Cas9 system to edit multiple loci in mammalian cells was demonstrated in 2013 (4, 6), numerous studies have demonstrated the multiplexing capacity of Cas9 system (63). Using mixes of RNP-guide RNA targeting, we have achieved co-targeting of two different loci, which may be a valuable tool to model tumoral heterogeneity in EC. As with most malignancies, EC arises from the accumulation of driver mutations that confer the hallmarks of cancer cell phenotype whilst contributing to generate heterogenous tumors, in which clones of tumoral cells with different mutational burden co-exist in a tumoral environment. Clones harboring most advantageous combinations of driver mutations will lead cancer progression. (64). Whole genome sequencing has unveiled or confirmed previously known mutations in EC (27). However, functional validation of alterations identified as a driver mutation itself or in combination with other alterations is still a challenge in cancer research. The main approach for functional validation of molecular alterations uncovered by whole genome sequencing of cancer is their recapitulation in genetically modified animal models, especially in mice. Traditionally, modelling of EC has been achieved using embryonic genetically mouse models carrying mutations in candidate genes. Such approaches provided unfading information about the role of genes alterations in EC development and progression. However, one of the main bottlenecks of this approach is the generation of mouse models carrying endometrial-specific mutations in more than one gene (i.e., double or triple knock-out) which requires breeding two or more genetically modified mouse strains with appropriated mouse carrying tissue specific Cre recombinase. These procedures may take even years to reach the desired combinations of mutant alleles to study their function in EC. Our *in vivo* CRISPR/Cas9-mediated genome engineering of mouse endometrium allows co-targeting of different loci in one single day, dramatically reducing the time to know the function of edited genes in EC. Another advantage is the temporal control and cellular specificity of genome edition. The availability of Cre-expressing mice that allow specific and inducible recombination in endometrial epithelial cells is still limited. In fact, there is no existing model that allows temporally controllable deletion of the genes of interest specifically in endometrial epithelial cells without affecting other cells into the uterus or other organs. By simply choosing the age of mice to be electroporated with the Cas9 RNP complexes, we can induce genome edition at different point of mouse lifetime. Since the incidence of cancers increases dramatically with age, temporal control of CRISPR/Cas9 delivery is an interesting feature of this method.

Another important hallmark of cancer is its monoclonal origin. Although the accumulation of genetic alteration created heterogeneity, all tumor cells arise from a single cell that received the initial mutation. However, most of conditional Cre-*LoxP* systems used to knock-out genes involved in EC, cause the ablation of the floxed gene in all cells expressing the Cre recombinase. Even Cre expressing mouse models that allow specific deletion of genes in endometrial epithelial cells genes deletion takes place in all epithelial cells, leaving no wild type epithelial endometrial cells in the whole uterus. In contrast, CRISPR/Cas9 mediated edition of mouse endometrial epithelial cells takes place in individual cells, resulting in a mosaic of edited cells side-by-side with wild type cells, which provides a closer scenario to tumoral microenvironment composition. Here, because of its well-established effects in mouse uterus, we used *Pten* as proof of concept of CRISPR/Cas9-mediated edition of endometrial epithelial cells. Nonetheless, we have demonstrated that other frequently mutated genes in EC such as *p53* or *Fbxw7* can also be edited. Future research using this technique will provide valuable functional information about the role of mutations in these and other candidate genes alone or in combination in the development of EC.

One of the possible pitfalls of the method is the failure to achieve biallelic ablation of targeted genes. To this regard, we have demonstrated the occurrence of biallelic edition of *Pten* gene that leads to a complete loss of PTEN protein expression and subsequent development of EC in a relatively short period of time. Nevertheless, we cannot rule out the possibility of monoallelic edition of one of the two copies of the targeted gene. Most probably, cells with monoallelic and biallelic edition of the targeted gene co-exist. In terms of a carcinogenic process, we think that this feature, far from being considered a drawback, may provide a good scenario to study haploinsufficiency requirements of tumor suppressor genes that further contribute to model tumoral heterogeneity. Besides the knowledge of gene functions in carcinogenesis, genetically modified models provide a platform for pre-clinical testing of anti-cancer drugs. Theus, disruption of a single gene or a combination of driver genes can generate new models to test the efficacy of targeted therapies.

Another important point is the reporting activity provided by the mT/mG dual fluorescent mouse. By achieving co-edition of the gene of interest (such as *Pten*) and the *LoxP* sites of the mT/mG locus, green cells carrying mutations in the gene of interest can be easily visualized and tracked by fluorescent microscopy techniques. Among them, Light Sheet Fluorescence Microscopy (LSFM) is an evolving optical imaging system for whole-volume imaging spanning cell biology, embryology, and *in vivo* live imaging. LSFM provides a high three-dimensional spatial resolution, high signal-to-noise ratio, fast imaging acquisition rate, and low levels of phototoxic and photodamage effects (65). The traditional method for histopathological analysis of samples involves a tedious and time-consuming process of dehydration, paraffin-embedding, sectioning and staining of samples. After this process a slide containing a section (or a few sections corresponding to different areas) of the tissue is obtained and microscopically analysed. Due to tumoral heterogeneity the observed sections from a sample may be not representative of the whole tumour structure and morphological properties and volumetric assessment of tumoral growth is not possible. LSFM imaging provides a comprehensive 3D overview of entire tissues and is complementary to conventional histopathological procedures (66), and in the case of heterogeneously distributed features in the tissue LSFM can become essential to fully characterize it. In this context, and as we have shown, LSFM imaging with efficient clearing techniques that preserve fluorescent proteins emission levels, may become a useful imaging tool for addressing three-dimensional volumetric growth of cancers in animal models and human biopsies in the near future (67). We have obtained volumetric imaging from mT/mG reporter mouse endometrium edited the *in vivo* CRISPR/Cas9 electroporation method, indicating that LSFM can be a useful microscopy technique to accurately examine tumoral growth in a three-dimensional perspective, allowing a precise modelling and assessment morphological features of EC. We believe that this method can be widely used to follow-up edited cell fate not only in the uterine cavity, but in other tissues where somatic CRISPR/Cas9 editing is intended.

In summary, we describe a new CRISPR/Cas9-based protocol for easy, efficient, robust, reproducible, rapid and flexible somatic genomic edition of mouse endometrial epithelial cells by RNP electroporation. We have validated this methodology as a faithful method for EC modelling. Nonetheless, we think that application of *in vivo* edition of mouse endometrial cells is not limited to endometrial oncology, but can also be applied to interrogate candidate genes for its role in the regulation of other important aspects of the uterine physiology, such as reproduction and development.

## AUTHOR CONTRIBUTION

Conceptualization, RN and X.D.; methodology, RN, ARM, MVS, APG, CML, LB, JC, DLN and X.D.; investigation, RN, ARM, MVS; validation, all authors; formal analysis, JE, ME, XMG, XD; resources, JC, LB, XD.; writing, RN, JC, XD.; supervision, XD.; main project administration, XD.; funding acquisition, XD.

## ACKNOWLEDGEMENTS

Supported by grants and PID2019-104734RB-I00 from Spanish Ministerio de Ciencia, Innovación y Universidades, Grupos estables de la Asociación Española Contra el Cancer (AECC) and LABAE19004LLOB. This work was also unded by Instituto de Salud Carlos III (MS17/00063) (co-founded by the European Social Fund (ESF), “investing in your future”). We wish to thank Sébastien Tosi from the Advanced Digital Microscopy core facility (IRB Barcelona) for support with image analysis tools.

## DATA AVAILABILITY STATEMENT

Data Availability Statement: The datasets analyzed during the current study are available from the corresponding author upon reasonable request.

