## Supplementary Methods for "Transient and DNA-free *in vivo* CRISPR/Cas9 genome edition for flexible modelling of endometrial carcinogenesis"

**Table S1**

| Targeted locus | crRNA sequence (without PAM) |
| --- | --- |
| *LoxP (mTmG plasmid)* | GUAUGCUAUACGAAGUUAUU |
| *Pten (Exon 5)* | AAUUCACUGUAAAGCUGGAA |
| *p53 (Exon 7)* | GGAGUCUUCCAGUGUGAUGA |
| *Fbxw7 (Exon 3)* | GUUGUUGGUGUUGCUCAACA |

**Table S3.**

| Target gene | Primer sequences (0.8 µM/rxn) | | PCR Protocol | | | Amplicon size |
| --- | --- | --- | --- | --- | --- | --- |
|  |  |  | **T (ºC)** | **Time** | **Cycles** |  |
| *Pten* | Fwd  Rev | GAGCACAATGCTTGTTTCCATG  TACACTAAGTGGATACAACCAC | 95 ºC | 2’ | 1 | 622bp |
|  |  |  | 95 ºC | 30’’ | 45 |  |
|  |  |  | 55 ºC | 30’’ |  |  |
|  |  |  | 72 ºC | 1’20’’ |  |  |
|  |  |  | 72 ºC | 7’ | 1 |  |
| *p53* | Fwd  Rev | CCATGCTAAGCAAGTGTTGG  CCCTAAGCCCAAGAGGAAAC | 95 ºC | 2’ | 1 | 451bp |
|  |  |  | 95 ºC | 30’’ | 45 |  |
|  |  |  | 55 ºC | 30’’ |  |  |
|  |  |  | 72 ºC | 1’20’’ |  |  |
|  |  |  | 72 ºC | 7’ | 1 |  |
| *Fbxw7* | Fwd  Rev | TTAAATACATCTGGGGCAAGC  TCCGCTTACATCTGGCTTCT | 95 ºC | 2’ | 1 | 427bp |
|  |  |  | 95 ºC | 30’’ | 45 |  |
|  |  |  | 55 ºC | 30’’ |  |  |
|  |  |  | 72 ºC | 1’20’’ |  |  |
|  |  |  | 72 ºC | 7’ | 1 |  |

**PCR protocol to generate amplicons for in vitro CRISPR/Cas9 nuclease activity.** DNA fragments flanking the target sequence of the RNPs targeting the indicated genes were amplified with the indicated primers and PCR conditions in 50µl PCR reactions using Taq polymserase (Biolools). fwd; forward. Rev; reverse.

**Table S3.**

| Gene | Primers (0.8 µM/rxn) | | PCR protocol | | |
| --- | --- | --- | --- | --- | --- |
|  |  |  | **T (ºC)** | **Time** | **Cycle** |
| *Pten* | Fwd  Rev | TTATCTTTTTACCACAGTTGCAC  TACACTAAGTGGATACAACCAC | 95 ºC | 2’ | 1 |
|  |  |  | 95 ºC | 30’’ | 45 |
|  |  |  | 55 ºC | 30’’ |  |
|  |  |  | 72 ºC | 1’20’’ |  |
|  |  |  | 72 ºC | 7’ | 1 |
| *p53* | Fwd  Rev | CCATGCTAAGCAAGTGTTGG  CCCTAAGCCCAAGAGGAAAC | 95 ºC | 2’ | 1 |
|  |  |  | 95 ºC | 30’’ | 45 |
|  |  |  | 55 ºC | 30’’ |  |
|  |  |  | 72 ºC | 1’20’’ |  |
|  |  |  | 72 ºC | 7’ | 1 |
| *Fbxw7* | Fwd  Rev | TTAAATACATCTGGGGCAAGC  TCCGCTTACATCTGGCTTCT | 95 ºC | 2’ | 1 |
|  |  |  | 95 ºC | 30’’ | 45 |
|  |  |  | 55 ºC | 30’’ |  |
|  |  |  | 72 ºC | 1’20’’ |  |
|  |  |  | 72 ºC | 7’ | 1 |
| *LoxP* | Fwd  Rev | TGGTTATTGTGCTGTCTCATC  GTGAGCAAGGGCGAGGAGCT | 95 ºC | 2’ | 1 |
|  |  |  | 95 ºC | 30’’ | 45 |
|  |  |  | 55 ºC | 30’’ |  |
|  |  |  | 72 ºC | 1’20’’ |  |
|  |  |  | 72 ºC | 7’ | 1 |

**PCR protocol to generate amplicons for NGS.** DNA fragments flanking the target sequence of the RNPs targeting the indicated genes were amplified with the indicated primers and PCR conditions in 50µl PCR reactions using Taq polymserase (Biolools). fwd; forward. Rev; reverse.

**Table S4**

| Antibody | Dilution | Source | Catalog | EnVision™ FLEX | Secondary antibody |
| --- | --- | --- | --- | --- | --- |
| PTEN | 1:100 | Dako | M3627 | *High pH* | EV FLEX Kit |
| Cytokeratin 8 | 1:200 | DSHB | AB531826 | *High pH* | Rat anti-biotin |
| GFP | 1:100 | Rockland | 600-101-215 | *High pH* | Goat anti-biotin |
| p-AKT (Ser473) | 1:50 | Cell signalling | 3787 | *High pH* | Rabbit anti-biotin |
| EnVision FLEX detection kit | RTU | Dako | K8002 | - | - |
| Goat anti-biotin | 1:200 | Santacruz | SC-2489 | - | - |
| Rat anti-biotin | 1:200 | ABCAM | AB6733 | - | - |
| Rabbit anti-biotin | 1:200 | Jackson | 111-065-144 | - | - |
| Estreptavidin-HRP | 1:400 | Dako | P0397 | - | - |

**Antibodies and immunohistochemistry conditions.** List of antibodies, dilution used, source and catalog number, antigen retrieval buffer and secondary antibody for each primary antibody are specified.
